# Retracing the Hawaiian silversword radiation despite phylogenetic, biogeographic, and paleogeographic uncertainty

**DOI:** 10.1101/301887

**Authors:** Michael J. Landis, William A. Freyman, Bruce. G. Baldwin

## Abstract

The Hawaiian silversword alliance (Asteraceae) is an iconic adaptive radiation of 33 species. However, like many island plant lineages, no fossils have been assigned to the clade. As a result, the clade’s age and diversification rate are not known precisely, making it difficult to test biogeographic hypotheses about the radiation. In lieu of fossils, paleogeographically structured biogeographic processes may inform species divergence times; for example, an island must first exist for a clade to radiate upon it. We date the silversword clade and test biogeographic hypotheses about its radiation across the Hawaiian Archipelago by modeling interactions between species relationships, molecular evolution, biogeographic scenarios, divergence times, and island origination times using the Bayesian phylogenetic framework, RevBayes. The ancestor of living silverswords most likely colonized the modern Hawaiian Islands once from the mainland approximately 5.1 Ma, with early surviving silversword lineages first appearing approximately 3.5 Ma. In testing the progression rule of island biogeography, we found strong positive evidence of the dispersal process preferring old-to-young directionality, but strong negative evidence for speciation occurring on islands during their young growth phase. This work serves as a general example for how diversification studies benefit from incorporating biogeographic and paleogeographic components.

## 1 Introduction

From Darwin’s finches in the Galápagos to the Greater Antillean anoles to the Hawaiian silverswords, adaptive radiations in island systems provide biologists with rare and precious glimpses into how macroevolutionary processes behave (e.g. Blonder et al. 2016; Kamath and Losos 2017; Lamichhaney et al. 2016). Adaptive radiations in island systems are particularly valuable to researchers as natural experiments, where island communities serve as replicates for studying the repeatability of evolutionary outcomes through ecological adaptation (Baldwin and Robichaux 1995; Gillespie 2004; Losos 1992; Losos et al. 1998b; Mahler et al. 2013; Grant et al. 2004). One feature characterizing adaptive radiation is the expansion of ecological adaptations among closely related lineages as they encounter new regions of niche space, where the radiating clade is often enriched for disparity and diversity relative to background rates of evolution (Givnish 2015; Schluter 2000; Simpson 1944; Osborn 1902). Since interspecific competition, access to new resources, and other spatiotemporal factors drive radiations, timing matters when discussing adaptive radiations.

Yet another common feature of many insular adaptive radiations, such as those mentioned above, is that they appear to result from few—or even one—long-distance dispersal event(s) from a mainland source area into an island system (Poe et al. 2017; Sato et al. 2001; Silvertown et al. 2005; Baldwin et al. 1991). Under such circumstances, several confluent factors complicate efforts to date when radiating lineages diverged. For instance, terrestrial species that are generally suited to dispersing over vast bodies of water—including plants, arthropods, small lizards, birds, and bats–have relatively sparse representation in the fossil record (Allison and Bottjer 2011). Once the ancestral lineage of an incipient radiation has established itself in its new setting, fossils must be recovered from the island itself in order to time-calibrate the internal divergence events of the radiating clade. Fossil recovery rates for terrestrial organisms within islands, and particularly within ephemeral oceanic islands, is low, notwithstanding significant finds from special sites such as lava tubes, bogs, sand dunes, and limestone caves and sinkholes (e.g. Burney et al. 2001; Hotchkiss and Juvik 1999; Olson and James 1982). When fossils are available, assigning them to key divergence events within a radiation is not necessarily easy, in part due to convergences of traits under island syndromes (Carlquist 1974; Gillespie et al. 1997; Losos et al. 1998a), the reduction of synapomorphies among anatomical features that readily fossilize (Sansom et al. 2010), and the often exceptionally short internode distances between the first divergences of a radiation (Gavrilets and Losos 2009). This set of circumstances is tantalizing to biologists because many of the features that make adaptive radiations in island systems appealing for study simultaneously undermine efforts to determine the ages—and, thus, estimate the evolutionary rates—of radiating clades. Ultimately, less precision in dating of clades within an adaptive radiation results in a weaker understanding of the timing and sequence of key events that constitute the radiation itself.

The silversword alliance (Asteraceae) represents one such adaptive radiation (e.g. Judd et al. 2016). Members of the silversword alliance form an endemic Hawaiian clade of 33 species nested within a larger clade corresponding to subtribe Madiinae, the tarweeds (Baldwin and Wessa 2000). Excluding the silversword alliance, nearly all remaining tarweeds are adapted to the Mediterranean-like climate of the California Floristic Province of western North America (Baldwin 2003b; Raven and Axelrod 1978). The biogeographic disjunction and phylogenetic relationship between the silversword alliance and continental tarweeds implies at least one long-distance dispersal event from the American mainland into the Hawaiian Archipelago. But an understanding of exactly when the first tarweed(s) initially colonized the Hawaiian Islands and when the silverswords began to diversify has been hampered by a lack of known fossils of Madiinae.

In one of the earliest molecular divergence time estimation efforts, Baldwin and Sanderson (1998) estimated the maximum crown age of the silversword alliance to be 5.2 (±0.8) Ma, made possible by integrating diverse lines of evidence. Continental tarweeds are almost entirely adapted to summer-dry conditions that began to develop in western North America at Mid-Miocene, approximately 15 Ma (Baldwin 2014; Jacobs et al. 2004). If crown tarweeds began diversifying only after the onset of such summer-dry conditions, as Baldwin and Sanderson reasoned, then tarweeds are at most 15 million years old. Using a clock-like nuclear ribosomal ITS tree with an external calibration of 15 Ma, Baldwin and Sanderson were able to estimate the maximum silversword alliance crown age and thereby compute the expected minimum speciation rate under a pure-birth process. They also noted that their maximum age estimate for the silverswords of 5.2 ± 0.8 Ma is remarkably consistent with the minimum age estimate of Kaua‘i of 5.1 ± 0.2 Ma (as the island’s age was known nearly twenty years ago; Clague and Dalrymple 1987).

Hawaiian paleogeographic evidence, however, did not enter into their dating estimate. At the time, use of such data for estimating divergence times or clade ages was problematical for multiple reasons. First, the complex geological history of the Hawaiian Archipelago was less well understood; today, island ages are known more accurately and precisely, but still not perfectly (Clague and Sherrod 2014). Second, despite the fact that tarweeds inhabit both North America and the major groups of Hawaiian Islands is evidence that some number of biogeographic events must have occurred, the events themselves are unobserved in terms of timing and geographical context. Finally, the distribution of (unobserved) biogeographic events depends on a phylogenetic context, which is also unobservable and must be inferred. These sources of paleogeographic, biogeographic, and phylogenetic uncertainty exist whenever biogeography is used to time-calibrate a phylogeny, though to different degrees for different systems.

Accurately dated phylogenies are necessary to test empirical biogeographic hypotheses about island radiations. An example of such a hypothesis is the “progression rule” of island biogeography. First articulated for hotspot archipelagos by Funk and Wagner (1995), the progression rule states that clades tend to inhabit older islands first and disperse to younger islands in the order that the islands appear. Adherence to this rule depends largely on ecological factors, such as whether the lineage may thrive in the context of the newly encountered community (Shaw and Gillespie 2016). Another factor is that younger islands have been biogeographically accessible for shorter periods of time compared to older islands, thus enabling fewer opportunities for a dispersing lineage to establish itself there. A related hypothesis is what we call the “speciation corollary” of the island biogeography progression rule: clades tend to experience higher rates of speciation when islands are young. The idea is that unspecialized lineages colonize new islands and then rapidly specialize and diversify as they fill available niches. This pattern was proposed by Wagner et al. (1995) to explain the Hawaiian radiation of *Schiedea* and *Alsinidendron* (Caryophyllaceae) and is predicted under the general dynamic model of island biogeography (Whittaker et al. 2008). Early work testing the progression rule hypothesis relied on pattern biogeography, such as area cladograms, to test for the rule’s existence in a clade (Cowie and Holland 2008; Gillespie et al. 2008; Parent et al. 2008; Funk and Wagner 1995). To our knowledge the rule’s speciation corollary has not been explicitly tested in a phylogenetic framework since it requires incorporating the timing of both speciation and paleogeographic events. Here we jointly model the phylogenetic, biogeographic, and paleogeographic processes of the silversword radiation, enabling us to statistically test how well the silversword radiation obeyed the progression rule of island biogeography and its speciation corollary.

The primary goal of this study is to illuminate the major biogeographic and evolutionary events accompanying the radiation of silverswords throughout the Hawaiian Archipelago. Such understanding depends on the diversification times within the silversword alliance, and most critically among those dates, the age of the most recent common ancestor (MRCA) of living members of the clade. We estimate these unknown ages using the process-based biogeographic dating technique described in Landis (2017) under the Dispersal-(Local) Extinction-Cladogenesis (DEC) model of Ree et al. (2005) to generate time-heterogeneous transition probabilities. To accomplish this, we simultaneously fit our dataset to an ensemble of phylogenetic models—including diversification processes, time-stratified biogeographic processes, and processes of molecular evolution—whose complementary features induce time-calibrated node age estimates. Additionally, we adapted the uniformization method (Rodrigue et al. 2008) of stochastic mapping (Nielsen 2002) to operate on time-stratified biogeographic processes in order to understand how and when the silversword alliance ancestor(s) first colonized the Hawaiian Islands and when diversification of the crown group began. Using this framework, we also developed new statistical tests for the progression rule of island biogeography and its speciation corollary that are informed by the timing and nature of dispersal and speciation events throughout the archipelago. Finally, we discuss the potential use of process-based biogeographic dating methods when studying other island biogeographic systems, and how the method may be improved.

## 2 Methods

We estimated the timing and ordering of the silversword radiation using a fully Bayesian phylogenetic analysis. Central to our analysis was the premise that paleogeographic dynamics induce time-heterogeneous biogeographic transition probabilities (Ree and Smith 2008). For example, a dispersal event into an island has probability zero before that island formed and a non-zero probability afterwards. In a phylogenetic context, this means that the biogeographic rate of events and the geological timing of events are separately identifiable. That said, the relative divergence times and topology of a phylogeny are not adequately estimated from biogeographic data alone. We concurrently estimate those aspects of phylogeny from molecular data (Zuckerkandl and Pauling 1962; Thorne et al. 1998). By combining sources of information from molecular, biogeographic, and paleogeographic evidence, our approach jointly models these features to estimate a geologically dated phylogeny. For more details on process-based biogeographic dating, see Landis (2017).

### Silverswords and tarweeds

We include 43 species and subspecies from the clade corresponding to tribe Madieae sensu Baldwin et al. (2002), including 35 taxa from the silversword alliance plus eight outgroup taxa. The silversword alliance is composed of three genera: *Argyroxiphium*, *Dubautia*, and *Wilkesia.* For molecular data, we obtained the same 647 bp multiple sequence alignment of the nuclear ribosomal internal transcribed spacer region (ITS) as used in Baldwin and Sanderson (1998). This dataset was chosen because it is highly congruent with phylogenetic evidence from nuclear chromosomal rearrangements (Carr 2003; Carr and Kyhos 1986), in contrast to chloroplast DNA trees, which are highly incongruent with ITS and chromosomal structural data as a result of chloroplast capture (Baldwin 1997; Baldwin et al. 1990) (B. G. Baldwin and W. A. Freyman, unpubl. data). Use of the same dataset analyzed by Baldwin and Sanderson (1998) also facilitates a comparison of the performance of our methods with those of the most detailed previous study of the age and rate of diversification of the silversword alliance. Hawaiian species ranges were coded according to Wagner et al. (2005), with two exceptions that are mentioned in the next subsection.

### Model

#### Geographical areas and paleogeographical uncertainty

The Hawaiian Islands are a Pacific archipelago located far from any continental flora. Of particular interest, the islands form an extensive chain in the North Pacific, from southeast to northwest, in sequence from youngest to oldest. The strict ordering of island ages has resulted from the relationship between the volcanic Hawai‘i hotspot, which produces newborn islands during eruption, and the steady northwesterly drift of the Pacific Plate over the hotspot (Clague and Sherrod 2014). Although the difference in neighboring island ages is semiregular, on the order of one to two million years, no island’s age is known perfectly without error. One component of the error may be caused by estimation error in dating the rock formations—an error term that will likely diminish with advances in geological methods—but a second component of uncertainty emerges from the fact that the date of a formation only provides a minimum bound on the island age (i.e., an island whose oldest estimated surfacing date is, say, 5 Ma must be at least that old, but it could be older). Introducing further uncertainty, we are interested in the maximum age at which the island was habitable in order to influence the dispersal rate of species into the island. Each island was formed over several stages of volcanic activity, where biogeographically relevant features, such as habitability and rock volume, vary between stages. For this study, we only considered the island formation times. Figure 1 provides the island age ranges we adopted, following the dates proposed by Lim and Marshall (2017), which buffer the minimally observable age estimates presented by Clague and Sherrod (2014).

**Figure 1:**
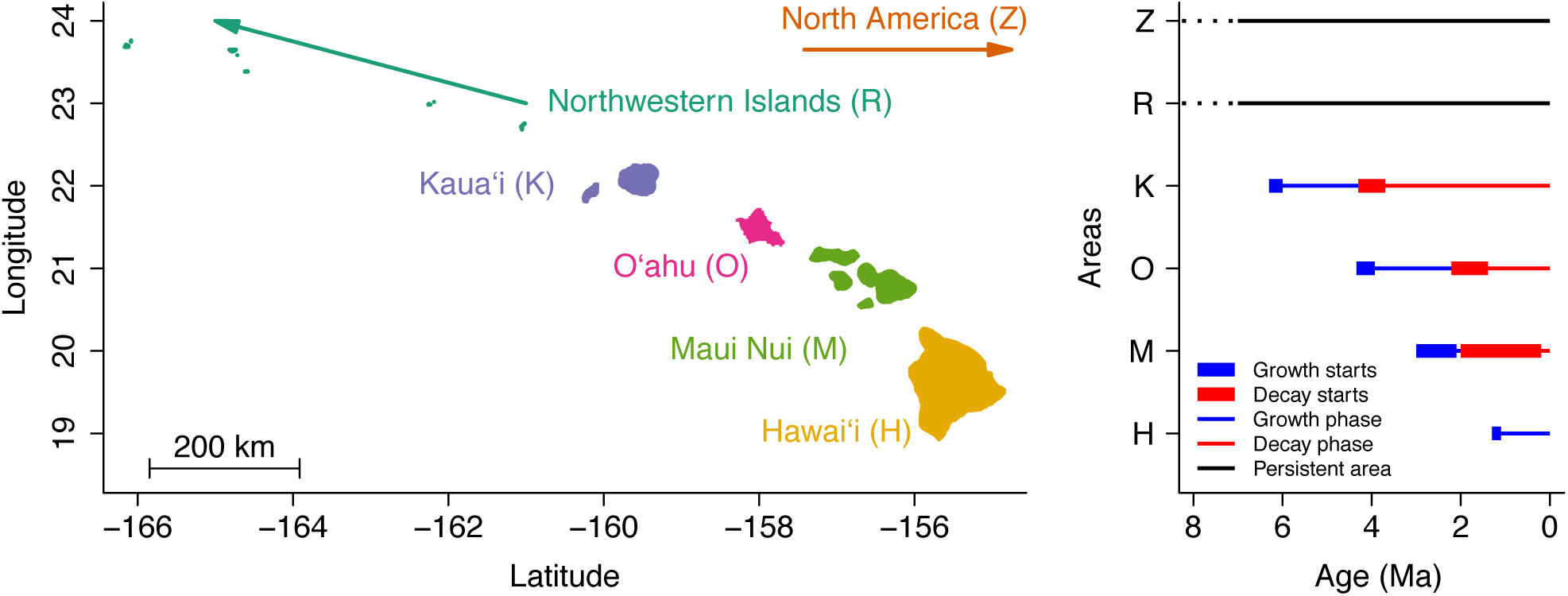
The Hawaiian Archipelago. The left panel shows the six biogeographic areas used in this study (in ascending order of age): Hawai‘i (H), the Maui Nui complex (M), O‘ahu (O), Kaua‘i and Ni‘ihau (K), the Northwestern Hawaiian Islands (R), and the North American mainland (Z). Each line in the right panel corresponds to the paleogeographic history of a particular area. Blue lines indicate when an island is growing and red indicates when an island is decaying. Thick lines indicate the range of ages during which the growth or decay phases began. Islands are only inhabitable after the growth phase begins. We do not consider the growth and decay phases for two areas, R and Z.

Because we are reliant on the island ages to inform our divergence time estimates, we must model this uncertainty in order to correctly propagate estimation error. To do so, we modeled the island ages as uniform random variables bounded by the ages provided in Figure 1. Our “relaxed rock” approach integrates over all combinations of tree topologies, divergence times, and island ages using MCMC, just as one integrates over divergence times and fossil taxon sampling times when applying the fossilized birth-death process (Heath et al. 2014).

The area Maui Nui (M) represents a complex of seven volcanic shields, and encompasses the four modern islands in the system, Maui, Moloka‘i, Lana‘i, and Kaho‘olawe. The Northwestern Hawaiian Islands (R), from Kure Atoll to Nihoa, are unified into a single complex of areas with a continuous presence of terrain above sea level for the past 27 Ma, well before the origin of the tarweed and silversword alliance clade studied here. The North American mainland (Z) is also sufficiently old to ignore uncertainty. We omitted the youngest island from the ranges of two taxa, *Dubautia laxa* subsp. *hirsuta* and *D. plantaginea* subsp. *plantaginea.* Reducing the state space improves the computational efficiency of the method, as described in Webb and Ree (2012). We omitted young islands because we expect that omitting old islands from ranges might cause some lineages to appear artificially young during inference.

#### Paleogeography-dependent range evolution

Long-distance dispersal events are rare relative to short-distance dispersal events as evidenced by estimates of colonization frequency of increasingly remote islands (Carlquist 1974). For the silversword alliance radiation, the distance between the North American mainland and the Hawaiian Islands is greater than the distances among the islands by more than an order of magnitude. This distance is a compelling reason to assume that the direct ancestor of living members of the silversword alliance colonized the Hawaiian Archipelago only once—something that seems exceedingly likely, but is not necessarily true. However, we do not know exactly the likelihood of such a long-distance dispersal event. With this in mind, we parameterized dispersal rates between islands to correspond to their relative coast-to-coast distances, meaning that the data inform the magnitude of the dispersal penalty (Webb and Ree 2012; Landis et al. 2013).

Because of the linear direction of island emplacement, we can assume that the relative distances between islands or island areas have remained essentially constant over time in terms of their sequential order (Carson and Clague 1995). Accurate paleogeographical distances between islands were not available, so we used modern distances between all islands for simplicity. Distances were measured coast-to-coast as the crow flies. To model the effect of distance on dispersal, we first define the relative distances between areas as *g_ij_*, which encodes the geographical distances between each area pair (*i*, *j*) divided by the mean distance over all area pairs.

Then we define the dispersal rates

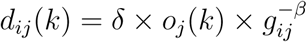

where *δ* is a base dispersal rate, *O_j_*(*k*) equals 1 if island *j* exists during epoch *k* and equals 0 otherwise, and *β* > 0 is a distance scaling parameter to be estimated. Note, the relative distance between any pair of areas equals 1 when *β* = 0.

By combining *d_ij_*(*k*), the dispersal rates, with *e*, the instantaneous extirpation rate, we construct the anagenetic dispersal-extirpation rate matrix, *Q*_DEC_(*k*), for each epoch *k* (Ree and Smith 2008). Ranges are constrained to be one or two areas in size to reduce the state space of the model (Webb and Ree 2012). Additionally, the original formulation of DEC results in extirpation rates that are biased towards zero when fitted to empirical datasets (Massana et al. 2015). Empirical datasets contain no extant taxa with size zero ranges (null ranges), resulting in ascertainment bias. To correct for this, we use conditioned transition probabilities,
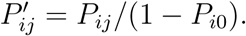

We model two types of cladogenetic events: allopatry, where each daughter inherits a mutually exclusive area that is a subset of the ancestral range; and sympatry, where one daughter lineage inherits the ancestral range while the other inherits just one ancestral area. Refer to Ree et al. (2005) for details.

Previous phylogenetic analyses demonstrated that the silversword alliance is securely nested within the primarily mainland North American tarweed clade (Baldwin et al. 1991) and allows us to constrain a North American (Z) root state. In effect, this requires that the mainland-island disjunction be explained by a dispersal into, rather than out of, the Hawaiian Islands. Combined, this lets us compute the likelihood of the range data given a phylogeny, range evolution parameters, and a (relaxed) paleogeographic hypothesis.

While we assume that a full-featured model most realistically portrays the biogeographic system, and thus favors more realistic evolutionary reconstructions, we would better understand which model features improve the results by contrasting such results to those of simpler models that are feature-poor. Two geography-aware models were considered: the full model allowed each modern island to appear independently in sequence (+G4), and the hotspot-naive model assumed that all four modern islands appeared in unison with the formation of Kaua‘i (+G1). As a point of contrast, we also considered a geography-naive model, which forced all areas to be present at all times and set the distances between all areas to be equal (–G). In essence, each model variant reconfigured how the dispersal rates between areas were computed. It is important to note that the -G model is time-homogeneous, so it contains no mechanism for the biogeographic process to inform the absolute timing of divergence events.

#### Molecular evolution and diversification processes

One point to emphasize is that we do not first infer the dated molecular phylogeny then subsequently model range evolution using the initial distribution of trees. Instead, range evolution is modeled simultaneously with the molecular evolution and diversification processes, allowing the biogeographic processes to inform the clade’s distribution of divergence times (Landis 2017).

Molecular variation is generated by a substitution process that treats the base frequencies and transition-transversion rate ratio as free parameters (Hasegawa et al. 1985). Site-rate heterogeneity is gamma-distributed (Yang et al. 1995a) with four rate categories and shape and scale priors uniformly distributed from 0 to 50. Branch-rate heterogeneity is modeled under an uncorrelated lognormal clock model (Drummond et al. 2006) with 32 discrete rate quantiles, where the mean clock rate has a uniform prior over orders of magnitude and the standard deviation is distributed by a exponential hyperprior with an expected value of one.

Diversification is modeled by a constant rate birth-death process (Nee et al. 1994), where the tree topology and divergence times are estimated as random variables. Recognizing that divergence time estimates are sensitive to modeling assumptions, we analyzed our data under a variety of birth and death rate priors, and a variety of taxon sampling scenarios. We took the birth and death rates to be exponentially distributed with mean values of 0.05, 0.10, 0.20, or 0.50. Assuming a uniform taxon sampling scheme, we considered three sampling probabilities of 0.17, 0.61, and 1.00, where the first value is an empirical estimate of the proportion of sampled species and subspecies within tribe Madieae, the second value is computed similarly but instead targets the *Madia* lineage (Baldwin 2003a), which contains the silversword alliance and closest mainland tarweed relatives (Baldwin and Wessa 2000), and the third value assumes perfect taxon sampling (see SI for details). Our presented results assume *Madia* lineage-wide sampling probabilities (*ρ* = 0.61) and moderate expected prior birth and death rates (0.10).

Three secondary node calibrations were generated by building upon recent work to date the radiation of Asteraceae (Barreda et al. 2015). We re-estimated the divergence times of Asteraceae using the original model settings of Barreda et al. (2015), but expanded the original backbone taxon set to include fourteen additional species belonging to and closely related to tribe Madieae, including the tarweed-silversword subtribe, Madiinae. Details for this analysis are provided in the SI. From the resulting posterior density, we translated the node age credible intervals (the 95% highest posterior density, or HPD95%) into node calibrations with uniform densities for the relevant nodes (Table 1). Pectinate backbone constraints were applied for the five oldest nodes in the phylogeny.

**Table 1:** Secondary node calibrations.

| Clade | mean age | minimum age | maximum age |
| --- | --- | --- | --- |
| Madiinae | 8.97 | 3.79 | 14.14 |
| *Arnica*+Madiinae | 10.75 | 4.91 | 16.59 |
| *Hulsea*+*Arnica*+Madiinae (root) | 15.99 | 9.13 | 22.85 |

### Analysis

#### Bayesian inference using RevBayes

This study relies on Markov chain Monte Carlo (MCMC) to estimate the joint posterior distribution of parameters for the molecular substitution process, the diversification process, the range evolution process, and paleogeographic features. All analyses were completed in `RevBayes` (Höhna et al. 2016). Additional analysis details are found in the SI. Analysis scripts and data files are available at github.com/mlandis/biogeo_silversword. Major features of the study are also available as teaching materials at revbayes.com/tutorials.

#### Secondary diversification rate estimate

The birth-death process used to model the diversification of all tarweeds and silverswords violates assumptions about uniform taxon sampling: it is fitted to a dataset that includes subspecies, and it assumes that the silversword alliance and mainland tarweeds diversified under the same rate-constant process. To obtain more empirically accurate rate estimates of species-level diversification, we estimated a second set of diversification rate parameters from the posterior distribution of silversword alliance diversification times. First, we applied a taxon filter to all trees in the posterior distribution used to construct Figure 2, pruning away all outgroup species and redundant silversword alliance subspecies to leave 25 of 33 known silversword alliance species. We then estimated the posterior birth-death process parameters from this set of species-level silversword alliance trees, treated as a mixture model over trees with uniform mixture weights. As priors, we assume the diversification rate is lognormally distributed and centered on two initial lineages giving rise to 33 species after the estimated silversword alliance crown age with log-standard deviation of 0.5 and a Beta(2, 2) prior on the turnover proportion.

**Figure 2:**
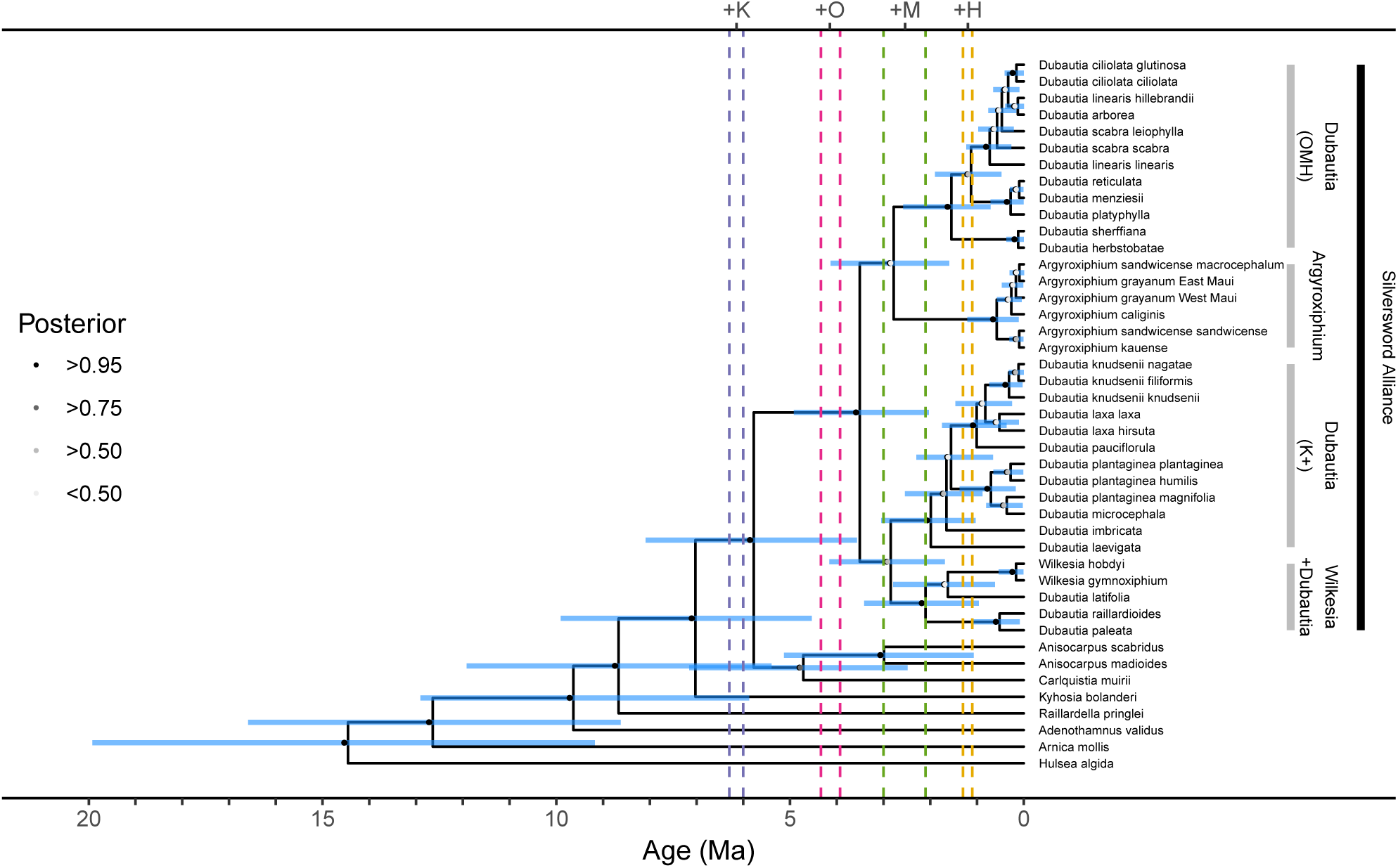
Maximum clade credibility tree of the silversword alliance and closely related tarweeds under the +G4 model. Vertical bars demarcate four subclades within the silversword alliance (see text). Node markers indicate posterior clade probabilities with shades from light gray to black. Node bars report the 95% highest posterior density for divergence time estimates. Vertical dashed lines bound the possible origination times per island complex (from left-to-right: Kaua‘i, O‘ahu, Maui Nui, Hawai‘i).

#### Stochastic mapping, ancestral state estimates, and summarizing uncertainty

We found that stochastic mapping by rejection sampling (Nielsen 2002) was inefficient for DEC, owing to the existence of an absorbing state (the null range), the asymmetry of rates, the large state space, and the underlying epoch model. For instance, rejection sampling will fail if the null range is sampled during a stochastic mapping, i.e., once a simulated history enters the null range, it remains in that state until the branch terminates, causing the sample to be rejected. This is almost certain to occur when the branch length is long or the extirpation rate is large. Particularly in the Bayesian setting, where parameters that do not maximize the likelihood are still of interest, stochastic mapping must perform reliably for all regions of parameter space with high posterior support. To address this problem, we modified the matrix uniformization sampling method described by Rodrigue et al. (2008) for the purposes of historical biogeography (Dupin et al. 2017). Our extension accounts for the time-stratified anagenetic and cladogenetic probabilities of DEC (see SI for details).

Ancestral range estimates were computed by sampling internal node states, before and after cladogenesis, using a demarginalization approach (Yang et al. 1995b). Ranges were sampled regularly during MCMC, and thus under a variety of evolutionary scenarios. To summarize the range estimates, using the maximum clade credibility tree as a reference, we omitted range samples corresponding to nodes whose left and right sister subclades were not found in the reference topology. For example, if the reference topology contained the node with subclades ((A,B,C),(D,E)), a sample containing the subclade (((A,B),C),(D,E)) would be a valid match, whereas a clade containing the subclade ((A,B),(C,(D,E))) would not.

#### Long-distance dispersal into the Hawaiian Islands

How did the ancestor(s) of the silversword alliance first colonize the modern Hawaiian Islands? The most parsimonious biogeographic scenario involves the direct colonization of the modern Hawaiian Islands, which must necessarily follow the formation of its oldest member, Kaua‘i. That said, less parsimonious scenarios are not strictly impossible. To measure the support of the probability of four categorical colonization scenarios: *single modern* involves one dispersal event directly from North America (Z) to a modern island (Kaua‘i, K; O‘ahu, O; Maui Nui, M; or Hawai‘i, H); *single older*-*single modern* describes one dispersal event to the older islands (R) then a second singular event to a modern island; *single older*-*multiple modern* is like the previous entry, but allows for multiple dispersal events from the older islands into the modern ones; and *multiple older/modern* requires more than one dispersal event from the mainland to any of the older or modern Hawaiian Islands. Support across scenarios was measured by querying the joint posterior of dispersal times, divergence times, and tree topologies, with each sample’s dispersal sequence being classified by a simple recursion.

#### Testing the progression rule of island biogeography

To test whether the silversword radiation obeyed the progression rule of island biogeography and its speciation corollary, we sampled stochastically mapped histories of the biogeographic process. As before, these samples incorporate all phylogenetic, biogeographic, and paleogeographic uncertainty defined by the model. For the progression rule of island biogeography, we label each dispersal event as a positive case if the newly colonized area is younger than its current island (e.g. *K* → *O*) and a negative case otherwise (e.g. *H* → *M*). Support for the speciation corollary is measured by the ratio of speciation events occuring on young islands (positive cases) versus on old islands (negative cases). We define an island as young while it grows until the point that it reaches its maximal area, and old after that threshold. Note, Hawai‘i is growing and considered young today. Applying this definition to the four modern islands, we partition each posterior island age sample into growth (young) and decay (old) phases by sampling from the “short” growth interval published by Lim and Marshall (2017). Taking the divergence time and ancestral range estimated for each node in a posterior sample, we classify each speciation event as young or old by the above criteria. As a concrete example, suppose that Kaua‘i originated at 6.2 Ma and its growth phase ended at 4.1 Ma. A speciation event on Kaua‘i at 5.0 Ma would be considered a positive case for the speciation corollary, while the same event at 2.1 Ma would be considered a negative event. Hidden speciation events that left no sampled descendants are not counted, which we expect will cause us to underestimate the number of older speciation events that occurred on now-old islands that were once young. If we find that the majority of our posterior density supports ratios of positive-to-negative events that are greater than one, we treat it as evidence in support for the progression rule and/or its speciation corollary.

## 3 Results

### Dating the silversword radiation

Our full-featured biogeographic dating analysis under the +G4 model recovers the silversword alliance as monophyletic (*p* = 1.00) and sister to the moderately supported clade (*p =* 0.78) formed by *Anisocarpus madioides*, *A. scabridus*, and *Carlquistia muirii* (Figure 2). Taxa within the silversword alliance fall into four highly supported clades (*p* > 0.99), all of which have crown ages that likely followed the formation of O‘ahu. Two of the four supported alliance clades inhabit Kaua‘i partly or exclusively: the clade containing two *Wilkesia* species plus three *Dubautia* species and one clade of only *Dubautia* taxa. The remaining two clades are composed of taxa found only among the younger islands of O‘ahu, Maui Nui, and Hawai‘i: the *Argyroxiphium* clade (not on O‘ahu) and a second clade composed entirely of *Dubautia* species. We find some support favoring a sister relationship between the two Kaua‘i-inhabiting clades (*p* = 0.62), but not enough to be certain of their exact relationship.

Assuming the +G4 model and the moderate diversification process configuration described above, the crown age of the silversword alliance is 3.5 Ma (95% highest posterior density, or HPD95%: 2.0 to 4.9 Ma). Under our refined diversification rate analysis, we estimate that the crown of the silversword alliance diversified at the mean rate of 1.07 species per lineage per million years (HPD95%: 0.20 to 1.91 spp/Myr).

Figure 3 presents the node age densities for five important silversword alliance clades: the silversword alliance crown group; the *Wilkesia* + *Dubautia* clade that is endemic to Kaua‘i; the *Dubautia* clade that is predominantly found on Kaua‘i (Dubautia K+); the *Dubautia* clade that is found only on O‘ahu, Maui Nui, and Hawai‘i (Dubautia OMH); and the *Argyroxiphium* clade. Although the +G1 and −G models misrepresent Hawaiian paleogeography, their results are useful for contrasting with the +G4 results. Under the geography-naive model (–G), the silversword alliance crown age is extremely responsive to the prior model settings, with the crown age appearing before or after the appearance of Kaua‘i. Conditioning on paleogeography (+G4 or +G1) greatly dampens how sensitive the silversword alliance crown age estimate is to model conditions.

**Figure 3:**
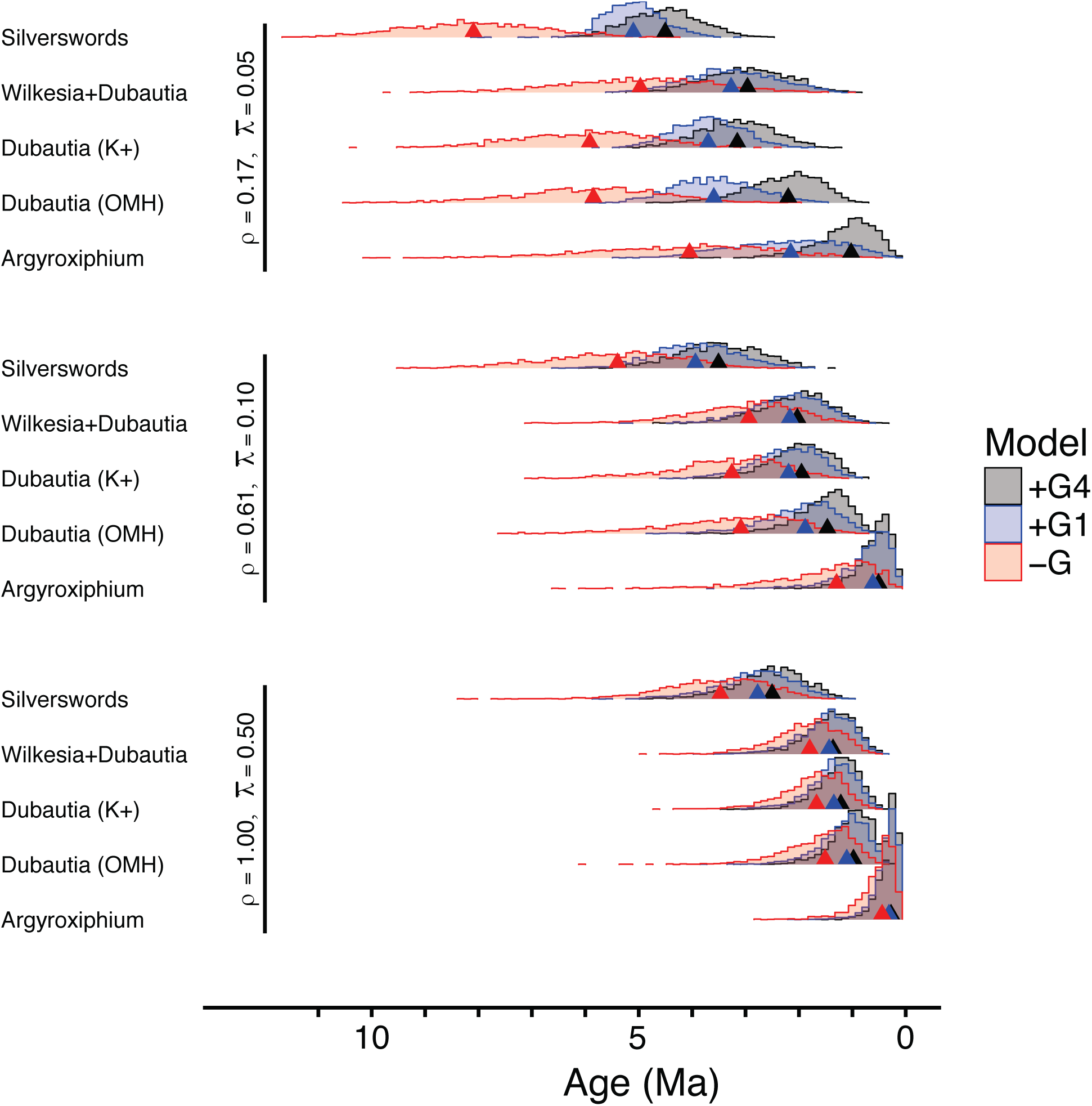
Silversword alliance clade ages under alternative model assumptions. Posterior clade age estimates for five highly supported clades and the three biogeographic models settings described in the text. The three panels in this figure show three of twelve diversification settings that were considered: slow birth/death rates and poor taxon sampling (top); moderate birth/death rates and medium taxon sampling (middle); high birth/death rates and perfect taxon sampling (bottom).

Examining the ages of the four highly supported silversword alliance subclades, we found that modeling the individual appearances of each island (+G4) generates additional dating information that is sacrificed when assuming all modern islands appear simultaneously (+G1). This effect is most evident when the diversification model assumes low sampling probabilities and slow prior birth and death rates (Figure 3, top); While *Argyroxiphium* and *Dubautia* (OMH) only inhabit modern islands younger than Kaua‘i, these two clades’ ages are frequently older than their island ages under the +G1 model, but not under the +G4 model.

Results for the remaining sensitivity analyses are given in the SI rather than here. However, one noteworthy result is that the divergence times are most consistent across subclades and model settings when we assume perfect taxon sampling and birth and death rate priors that favor exceptionally high birth and death rates (0.50). These settings induce a “tippier” tree, where all divergence times become sufficiently young that island availability no longer restricts dispersal patterns.

### Long-distance dispersal into the Hawaiian Islands

Figure 4 shows that under the fully featured +G4 model, the “single modern” scenario is favored to explain how tarweed ancestors first colonized the modern Hawaiian Islands (the +G4 probabilities of Figs. 4A–D sum to *p* = 0.87). Together, colonization scenarios involving a single long-distance dispersal event into the Hawaiian Islands (Fig. 4A–F) are roughly 13 times as probable as scenarios involving multiple events (Fig. 4G). When ignoring geography under the –G model, we find increased support for the “multiple older/modern” scenario, decreased support for either of the two “single older” scenarios, and decreased support for Kaua‘i as the destination under the “single modern” scenario. Estimates under the single-island +G1 model capture features of both the +G4 and –G analyses: +G1 is more similar to +G4 in that a single long-distance dispersal event is strongly favored, but more similar to –G in that support for Maui Nui as the destination is substantially increased relative to that for Kaua‘i. Lastly, the two “single older” scenarios find the greatest support under the +G4 model, indicating that support for the indirect colonization of the modern Hawaiian Islands may not be independent of the ages at which younger islands appear (+G1).

**Figure 4:**
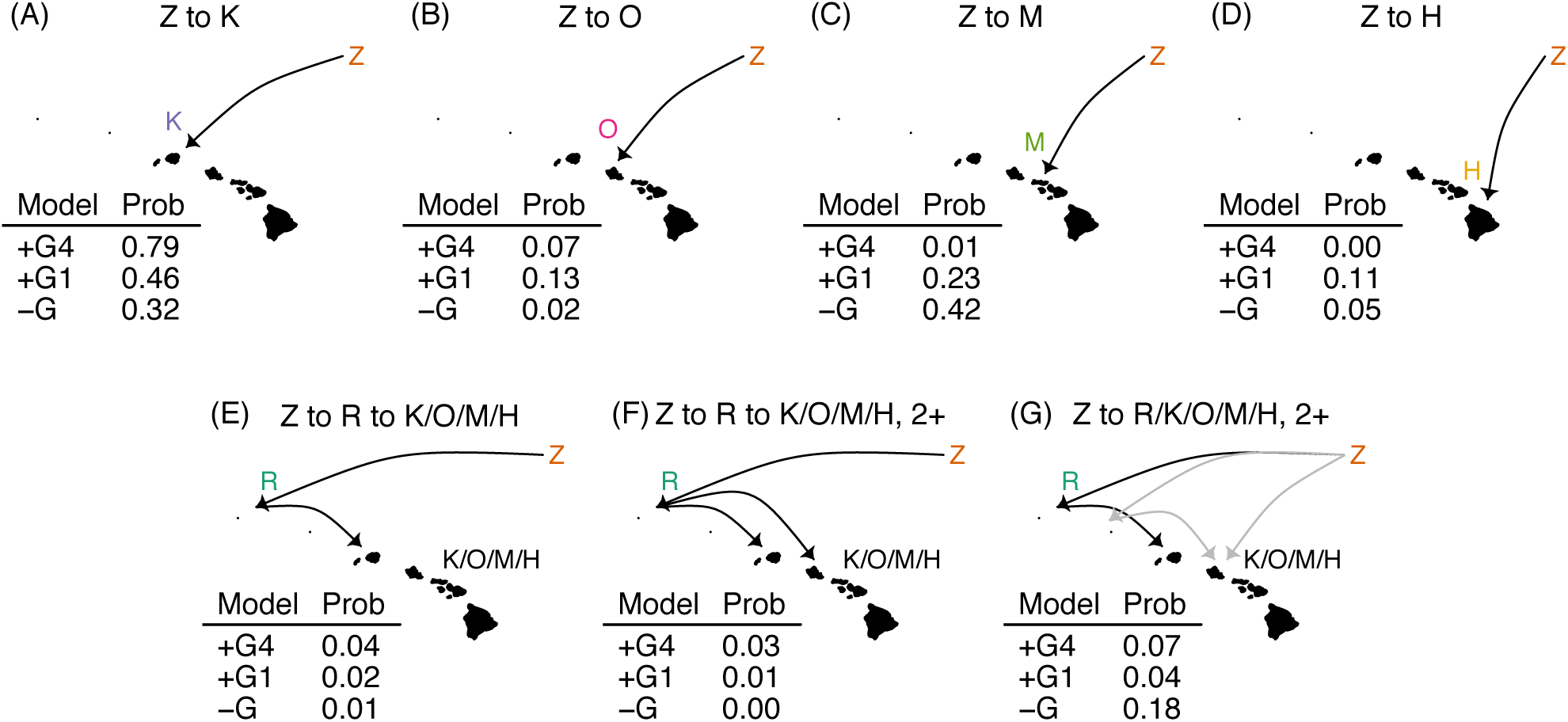
Hawaiian silversword colonization scenarios. Biogeographic dispersal histories were classified into several distinct evolutionary scenarios as described in the main text: (A-D) the single modern colonization scenario is represented with each of the four modern islands as destinations (panels A, B, C, D, correspond to K, O, M, H); (E) the single older-single modern scenario without regard to which modern island was the destination (K/O/M/H); (F) the single older-multiple modern scenario; and (G) the multiple older/modern scenario. Each scenariO‘s posterior probability is given for the three models: the full model (+G4), the hotspot-naive model (+G1), and the geography-naive model (–G).

### Dating key biogeographic events in the Hawaiian radiation

Figure 5 summarizes the joint distribution of phylogenetic and ancestral range estimates under the +G4 model as previously described in the Methods section. Consistent with the results presented in Figure 4, the ancestral range of the silversword alliance crown group very probably included the island of Kaua‘i. The majority of biogeographic variation appears in two clades: the *Argyroxiphium* clade and the *Dubautia* clade containing taxa on islands younger than Kaua‘i (i.e. the minimal clade including *D. arborea* and *D. sherffiana).* In the second major *Dubautia* clade, made largely of taxa that are endemic to Kaua‘i, the three dispersal events from Kaua‘i into the younger islands are relatively recent, occurring within the past one million years.

**Figure 5:**
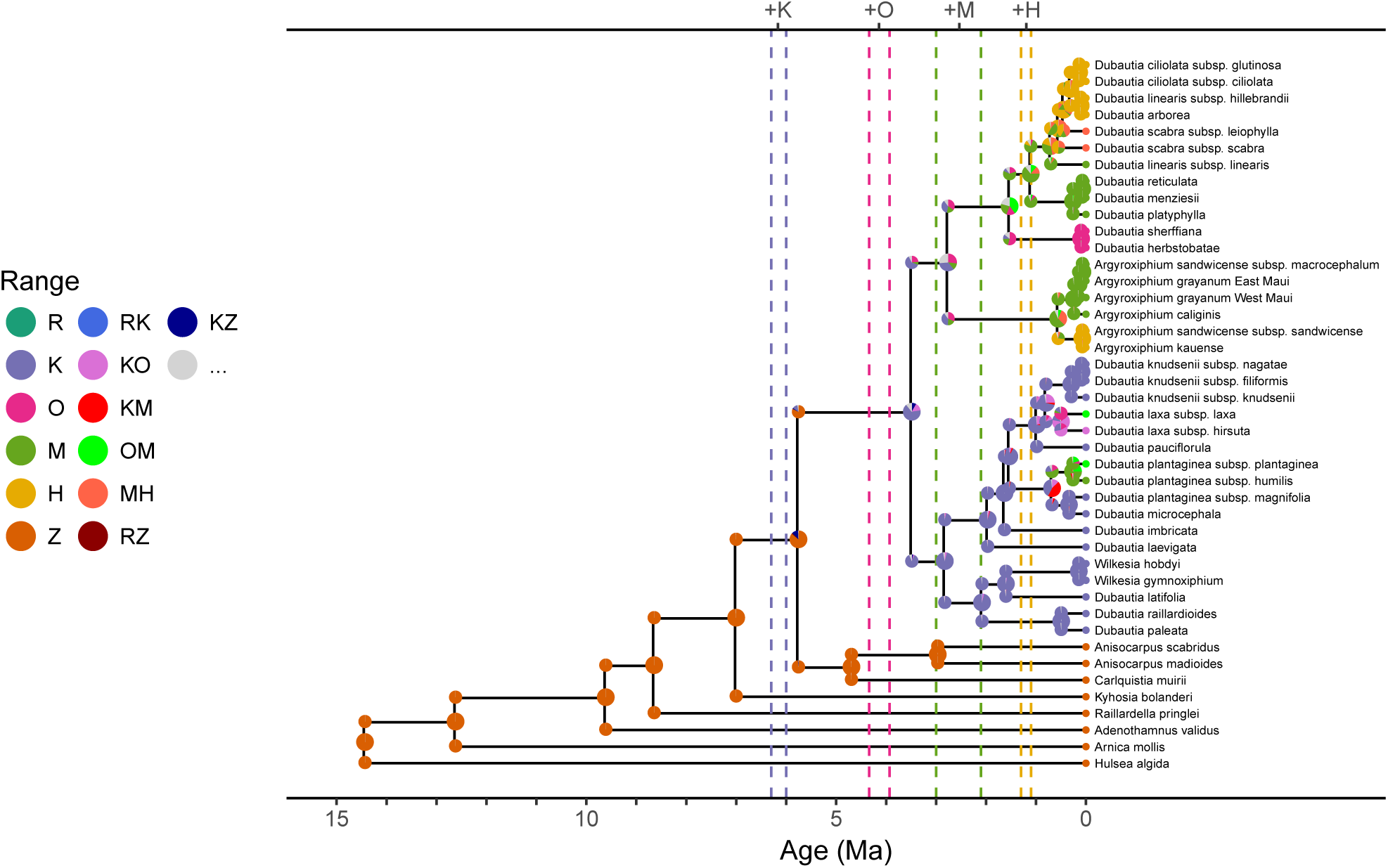
Ancestral range estimates of tarweed+silverswords under the +G4 model. Ancestral range estimates are plotted upon the maximum clade credibility tree. Pie charts report the range estimate probabilities before cladogenesis (node) and after cladogenesis (shoulders). The three most probable ranges are plotted per node/shoulder, with the remaining less probable ranges being binned into the range labeled ‘…’; these improbable ranges are valid in the model but not listed in the legend. Vertical dashed lines bound the possible origination times per island complex (from left-to-right: Kaua‘i, O‘ahu, Maui Nui, Hawai‘i).

The ancestral range estimate summary shown in Figure 5 does not display precisely when a particular island was first colonized nor report how those times might vary in response to phylogenetic uncertainty. To disentangle when key clades originated, when islands originated, and when those islands were first colonized, we present the posterior event ages in Figure 6A.

**Figure 6:**
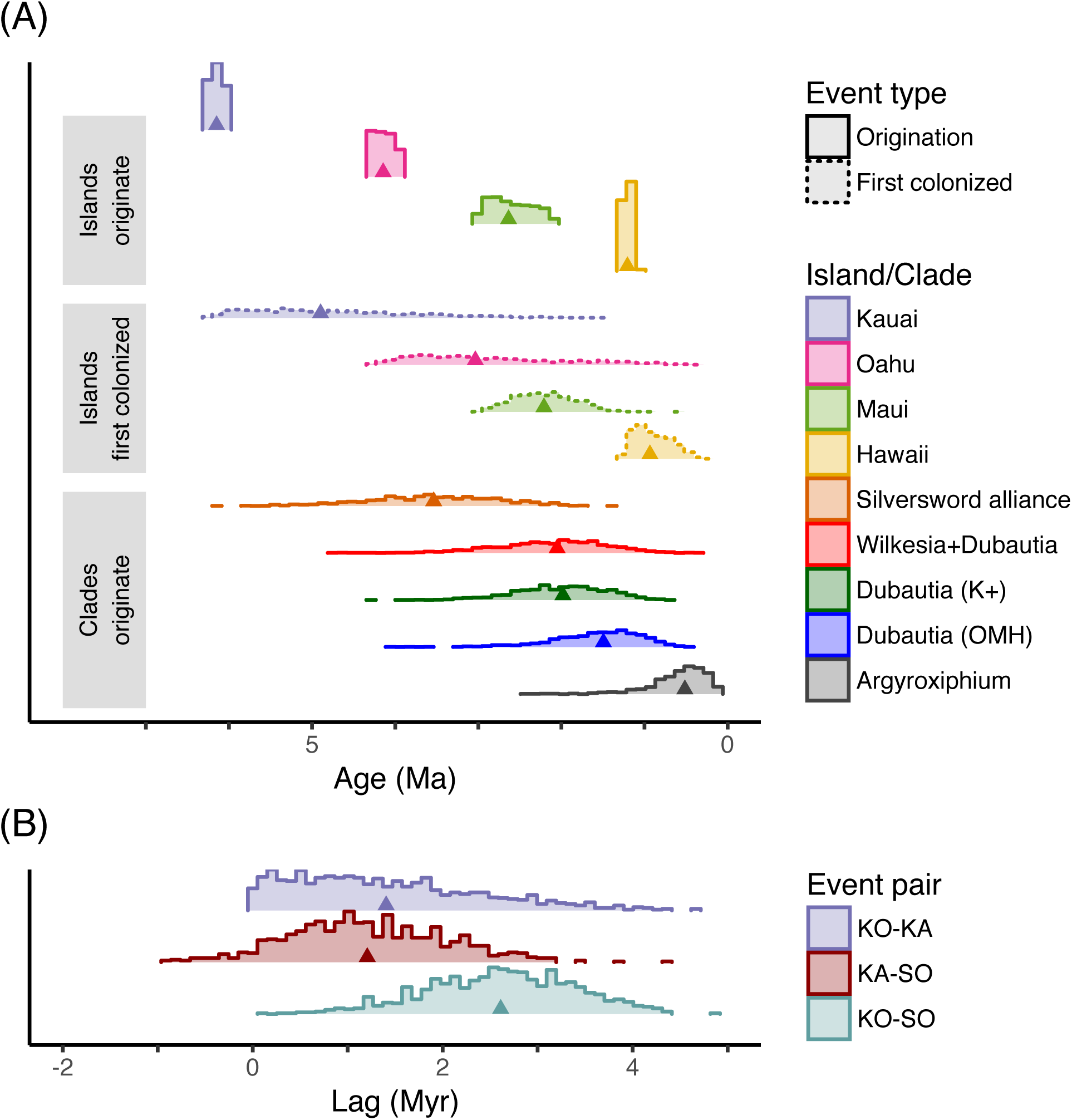
Age distributions of key biogeographic events in the silversword alliance radiation. (A) Histograms with dotted boundaries refer to first arrival times, while solid boundaries refer to origination times of islands and focal clades in this analyses (distinct by color). First arrival times relate to the first time any crown silversword alliance lineage arrived on a given island. Clade ages match those given for the +G4 model in the middle panel of Figure 3. Note, first arrival events always follow the origination time of the corresponding island. (B) Histograms show the posterior differences in time for pairwise combinations of the following three event ages: origination of Kaua‘i (KO); the ancestors of living silversword alliance species first arrive at Kaua‘i (KA); and the crown age of surviving members of the silversword alliance (SO). Note, KO-KA is always greater than zero because Kaua‘i cannot be colonized before it originates. The remaining differences, KO-SO and KA-SO, may be negative if the silversword alliance began to diversify before Kaua‘i formed or before the first arrival on Kaua‘i, respectively. The posterior mean differences are KO-KA=1.4, KA-SO=1.2, KO-SO=2.6 in millions of years.

The silversword alliance radiation throughout the modern Hawaiian Islands must have been precipitated by three historical events: the modern islands must have begun to form, the ancestor of living members of the silversword alliance must have first colonized the modern island chain, and the oldest surviving silversword alliance lineages must have begun to diversify. Even if improbable, the origination of the silversword alliance could have predated the origination of, or their arrival upon, the modern islands. Moreover, there was likely some delay between these critical events from the standpoints of biology, based on observations in community assembly and the element of chance in dispersal dynamics, and of mathematics, because the expected waiting time between dispersal events is necessarily greater than zero. Figure 6B reports the evolutionary lag between these events, taking the oldest island complex, Kaua‘i, as an upper bound. The delay between the island origination time and the first arrival time upon Kaua‘i is nearly 1.5 million years (posterior mean KO-KA=1.4 Myr), and between the first arrival time and the silversword alliance origination is over another million years (posterior mean KA-SO=1.2 Myr). Note, the posterior of lag separating the arrival at Kaua‘i from the crown age of the silversword alliance contains left tails that are negative, which is corroborated by the results presented in Figure 4.

### Testing the progression rule in silverswords

The silversword alliance radiation presents strong positive support for the progression rule of island biogeography (*p* > 0.99) with a posterior mean of 3.9 positive dispersal events for every negative dispersal event (Figure 7A,C,E). Consistent with earlier findings from Figure 4, we find greater support for dispersal from the mainland (Z) to Kaua‘i (K) than to the older Hawaiian Islands (R) or the remaining young islands (O/M/H). There is strong negative support for the progression rule’s speciation corollary in the silversword alliance (*p* < 0.01), however, with 3.8 speciation events occurring on older decaying islands for every speciation event occurring on younger growing islands (Figure 7B,D,E). Negative speciation events occurred primarily within Kaua‘i (61%), followed by Maui Nui (32%) and O‘ahu (7%), while positive speciation events most frequently occurred within Hawai‘i (60%), then Maui Nui (21%), then Kaua‘i (10%), and then finally O‘ahu (9%).

**Figure 7:**
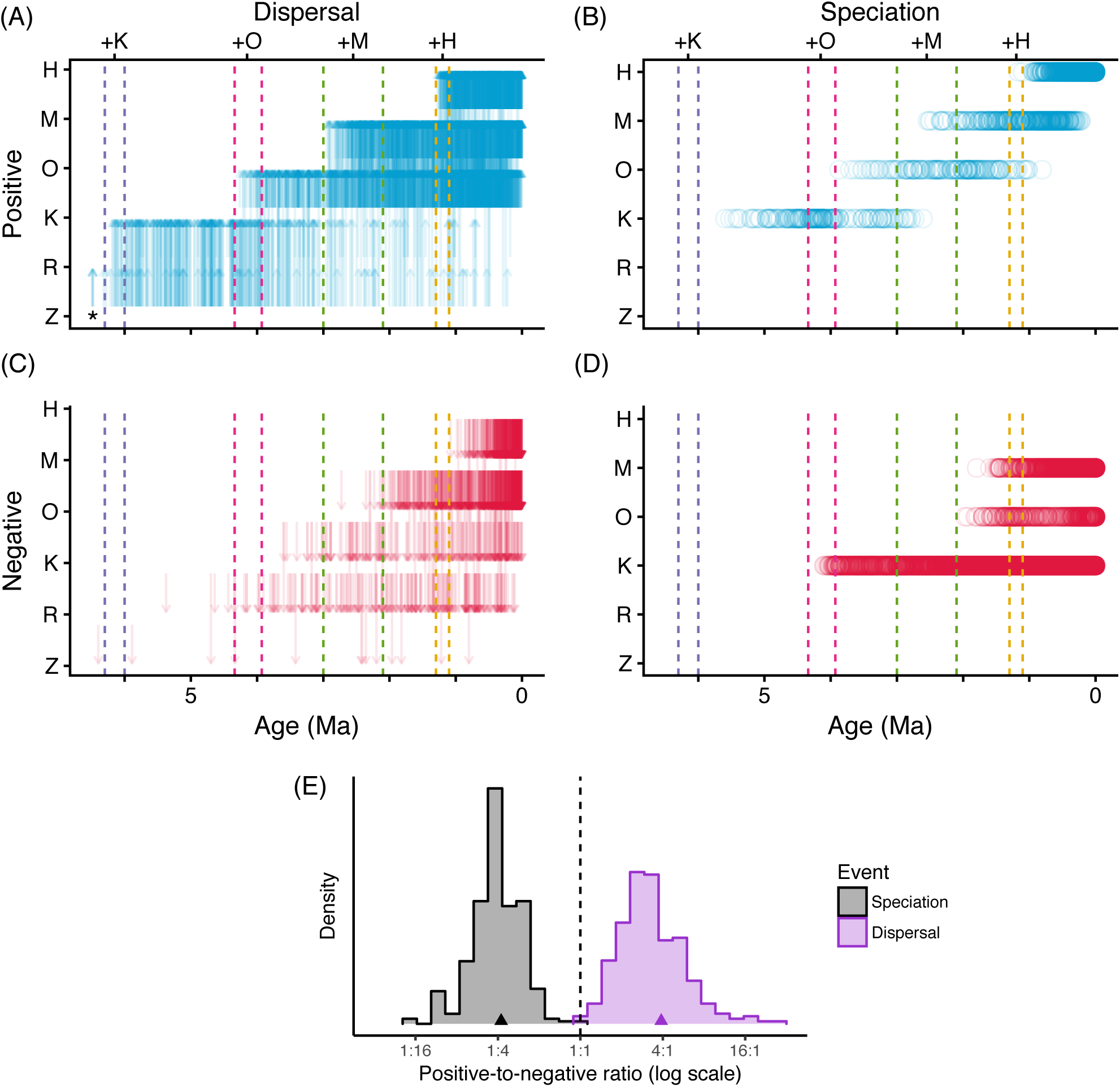
Posterior support for and against the progression rule of island biogeography and its speciation corollary. Dispersal events (A,C) and speciation events (B,D) are classified into either positive (A,B) or negative (C,D) cases that follow or break the progression rule. Positive speciation events occur on islands before the island reaches its maximal area (the older Hawaiian Islands, R, and the mainland, Z, are omitted). Positive dispersal events occur from an old area into a younger area. Dashed lines bound the possible origination times per island complex. (A) A small fraction of type-positive Z → R events occurred before 6.5 Ma, and are represented by the single arrow and asterisk. (E) Posterior estimates of the ratio of positive-to-negative cases of the progression rule for dispersal (purple) and the speciation corollary (black). Neither speciation nor dispersal processes are probable at the 1:1 ratio (dashed line); the dispersal process obeys the progression rule yet speciation events run opposite to the progression rule’s speciation corollary.

## 4 Discussion

The silversword alliance is an especially prominent example of insular adaptive radiation for which rigorous estimation of the timing of diversification and the pattern of inter-island dispersal have been long available (Baldwin and Robichaux 1995; Baldwin and Sanderson 1998). As such, the clade is ideal for examining the potential for advancing understanding of the radiation through a new approach to biogeographic hypothesis testing (Landis 2017). In particular, we used an integrative Bayesian phylogenetic framework incorporating refined paleogeographic information to disentangle colonization and diversification history and to test the progression rule of island biogeography (Funk and Wagner 1995) and its speciation corollary. Those findings provide a new perspective on the geological context of dispersal and evolutionary radiation in an insular clade, as discussed below.

The crown group of the silversword alliance began diversifying approximately 3.5 ± 1.5 Ma. This age is younger than 5.2 ± 0.8 Ma, the maximum clade age of the silversword alliance estimated by Baldwin and Sanderson (1998), whose results at the time provided early robust evidence for a major Hawaiian radiation that was contemporary with the modern high islands, rather than pre-dating the oldest high island, Kaua‘i. Where our analysis uses island paleogeography to date the clade, Baldwin and Sanderson’s maximum age estimate was obtained by applying a phylogenetic niche conservatism argument to paleoclimatological data from western North America in order to bound the maximum age of continental tarweeds. The consistency between the two age estimates suggests future work might integrate both lines of reasoning to further improve age estimates for tarweeds and the silversword alliance. The relatively tight support interval for the Baldwin and Sanderson estimate (±0.8 Ma) is smaller than ours for at least three reasons: we modeled the uncertainty in the root age estimate itself rather than the bound, our divergence time estimates assume rate heterogeneity across lineages rather than clock constancy, and Bayesian credible intervals and frequentist bootstrap replicates are not strictly equivalent.

As to the pace of the radiation, we estimate that the silversword alliance crown group diversified at a mean rate of 1.07 ±0.86 spp/Myr. Our diversification rate estimate differs in several ways from the minimum speciation rate estimate of Baldwin and Sanderson, 0.56 ± 0.17 spp/Myr. Part of the difference is explained by replacing the maximum age estimate (5.1 Ma) with an actual age estimate (3.5 Ma). The original estimate also assumed a pure birth process with perfect taxon sampling, and, technically, estimated the rate of new species or subspecies originating. Another difference is that we now obtain some internal node calibrations under the paleogeographic model, causing some lineages to appear at younger ages than they would if paleogeography was ignored (Figure 3).

Matching intuition, our analysis found it to be highly likely that the ancestor of living silverswords was a single tarweed lineage that first colonized Kaua‘i directly from the North American mainland, then began to diversify into what we know as the silversword alliance today. By ignoring the influence of geography on dispersal rates, multiple independent long-distance dispersal events rise in probability, from 0.07 to 0.18, suggesting a stronger influence of distance than island age in this result. Based on our analysis, it is unlikely that the Northwestern Hawaiian Islands, which arose prior to Kaua‘i, played a significant role in the tarweed colonization of the Hawaiian Archipelago. Modeling the effect of higher rates of extinction on those older islands, applying a slower prior to the diversification or substitution rates, or extrinsic information pushing the silversword alliance crown age to be older could overturn this result. But, as it is, nothing requires a colonization event into the Hawaiian Archipelago before the formation of Kaua‘i. Our estimate that both the crown *and* stem ages of the silversword alliance are contemporary with Kaua‘i corroborates the biogeographic importance of a pre-Kaua‘i gap in island formation and presence of only relatively small, widely spaced islands in the chain prior to the rise of Kaua‘i (Price and Clague 2002).

This lone long-distance dispersal event becomes an appealing candidate for use as a biogeographic node calibration, where one might assert that the silversword alliance crown group began to diversify only after Kaua‘i formed. Baldwin and Sanderson (1998) noted that any error in an island age estimate would result in a cascade of node age estimation error throughout the phylogeny, especially at deeper nodes. Supposing that the age of Kaua‘i was known perfectly, there is still the issue of what density to apply to the node: the density would need to measure the delay between the appearance of the island and the colonization of the island, and between the colonization of the island and the first speciation event that left sampled descendants. In other words, the biogeographic node age density should depend on the age of the island, the dispersal rate into the island, and the speciation rate on the island, but the values of those parameters are unknown and inferred through the evolutionary analysis itself. Sidestepping these complications by jointly inferring the evolutionary parameters along with the divergence times, we estimate this lag from the data directly rather than assert its effect through the prior (Figure 6B).

While the conspicuous disjunction between the continental tarweeds and the Hawaiian silversword alliance offers a singular plausible biogeographic event suitable for node calibration, weaker node calibrations could not be so easily or consistently applied to less certain biogeographic events within the silversword radiation. For example, six taxa of *Dubautia* sect. *Rail-Uardia* (the six taxa at far upper right of Fig. 2) are found on the youngest island, Hawai‘i. Suppose one was certain that these six taxa formed a clade. Their co-occurrence on Hawai‘i could be explained by a single dispersal event. The dispersal event must have occurred after Hawai‘i originated, thereby informing the age of the clade, which could justify the use of a biogeographic node calibration. But if we cannot be certain of the monophyly of the six taxa, then anywhere from one to six dispersal events may be needed to explain their occurrences, and the placement of those hypothetical events would need to be defined over the set of relevant clade topologies—and sets defining random treespace grow rapidly with the number of taxa. While node calibrations are not easily applied in cases such as this, process-based biogeographic dating methods inherently marginalize over all defined phylogenetic and biogeographic scenarios.

We found some effect for this subtler dating information when examining crown ages for the four major silversword alliance subclades. Three models were used: one model ignoring geography (–G), one model that reflects our best translation of paleogeography (+G4), and one model that ignored finescale paleogeographic knowledge (+G1). If the divergence times for the silversword alliance and the four subclades therein are equal when assuming +G1 or +G4, then the exact ages of appearance for O‘ahu, Maui Nui, and Hawai‘i would be inconsequential to dating the clade. However, we found that the ages estimated for the subclades that are endemic to the younger islands are older under +G1 than under +G4, indicating that fine scale phylogenetic, biogeographic, and paleogeographic interactions generate information that may be extracted through process-based biogeographic dating methods. This suggests that other datasets may contain similarly diffuse information about clade ages, a feature overlooked by traditional node calibration based frameworks.

Using the posterior distributions of speciation and dispersal events obtained from our stochastically mapped biogeographic histories, we classified the events as evidence in favor (old-to-young) or against (young-to-old) the progression rule and its speciation corollary, and we measured the probability and magnitude of support for and against the progression rule in island biogeography (Funk and Wagner 1995). While we found strong support (*p* > 0.99) for the dispersal process following the progression rule, the speciation process unequivocally does not follow the progression rule’s speciation corollary (*p* < 0.01). The negative signal for the speciation process is fueled by two of the silversword alliance subclades that have been diversifying without rest in Kaua‘i for millions of years. At a glance, the number of speciation events per unit time (i.e. the rate) remains remarkably steady within Kaua‘i, even as Kaua‘i declines in area. That finding cannot be explained by differences in taxonomic judgment about species boundaries in silversword alliance lineages on Kaua‘i versus the younger islands; species of the silversword alliance on Kaua‘i are in general even more divergent genetically than on younger islands (Carr and Kyhos 1986; Witter and Carr 1988) and are highly distinctive morphologically and ecologically (Carr 1985; 1999). Our result may instead reflect ecological factors associated with the extensive, deep erosional dissection of Kaua‘i that has accompanied its loss of area. To some extent, such activity may have offset habitat loss associated with reduction in island size by creating habitat heterogeneity (see Lim and Marshall 2017) and opportunities for isolation at finer geographic scales that have facilitated speciation, as predicted by (Whittaker et al. 2008) for islands at a comparable developmental stage (“maturity”) within oceanic archipelagos. Such considerations may be reflected by the relatively high number of silversword alliance species of limited geographic distribution on Kaua‘i (Carr 1999).

While we see the results of this process-based test of the progression rule as an advance in the study of island biogeography, it has limitations. One challenge arises in objectively defining when an island is young or old. This is simple for directional events, like dispersal, where for any pair of areas one is older than the other. Whether a speciation event occurs on a young or an old island is not so clear; we used the time when an island’s growth rate turned negative to demarcate young from old, but, as discussed above, other ecological factors may to some extent counteract loss of island size in influencing speciation rate. As another example, the ratio tests consider the proportions of positive and negative events within each stochastically mapped evolutionary history, then ask whether the ratio is generally larger, smaller, or equal to the balanced ratio of one-to-one. But those negative speciation events that took place in Kaua‘i over the past four million years may drown out evidence that some subclades positively adhere to the progression rule’s speciation corollary—namely the two subclades inhabiting only islands younger than Kaua‘i. Our analysis also did not consider possible extinction events on Kaua‘i (and elsewhere), where loss of higher elevation habitat through erosion and subsidence, for example, may have resulted in a bias toward loss of earlier diverging lineages. This bias might be eliminated by reformulating the DEC biogeography model within the State-Dependent Speciation and Extinction (SSE) framework (e.g. Goldberg et al. 2011). Such an approach could account for both cladogenetic events as well as extinction during the diversification process (Goldberg and Igić 2012; Freyman and Höhna 2017) and incorporate the effects of island ontogeny on speciation and extinction rates (Lim and Marshall 2017).

Considering the finer-scale features of our reconstruction, the novel finding of a considerable lag between island colonization and diversification of the silversword alliance (Figure 6B) is especially intriguing in light of the additional finding here of strong negative evidence for diversification during the island growth phase, at least on the oldest high island, Kaua‘i (Figure 7D). Although an undetected extinction bias toward early diverging lineages may partially explain these results, there remains strong evidence that diversification of the silversword alliance on Kaua‘i has continued apace as the island has diminished considerably in size through erosion and subsidence. The importance of new opportunities for speciation during the later stages of island development that may arise from such processes as erosional dissection of the terrain into more complex and isolated habitat space warrants more study and may help to explain why Kaua‘i contains higher species richness of the silversword alliance and of endemic angiosperms in general than any younger island of the chain (Sakai et al. 2002; Wagner et al. 2005).

Our work shows that a variety of biogeographic hypotheses may be tested by defining categorical hypotheses, then recording the frequency of events from the posterior distribution of stochastically mapped biogeographic histories (Dupin et al. 2017). Because our stochastic mappings are are fully Bayesian, they exactly characterize our confidence in the variety of biogeographic scenarios that are probable under the model. The fully Bayesian approach reports our uncertainty in both the biogeographic history *and* in the evolutionary and paleogeographic conditions that could have plausibly generated that history, thus guarding against a false sense of precision regarding past events. These estimates are subtly distinct from maximum likelihood settings where histories are typically simulated under the single point estimate of parameters with the highest probability, rather than over the range of model parameters with high probability (i.e. those with high confidence/credibility). That said, obtaining Bayesian stochastic mappings over a range of probable parameters posed some technical challenges. Rejection sampling approaches for stochastic mapping were, for all practical purposes, incompatible with posterior samples where the rate of area loss was not small and/or branch lengths were long; nearly all simulated stochastic mappings under the biogeographic process lead to the null range (an absorbing state) under these settings. To circumvent this issue, we extended the uniformization sampling method (Rodrigue et al. 2008) to accommodate cladogenetic events and the time-heterogeneous rate matrices of the epoch model.

Although some of the coarse-scale features of our reconstruction may not surprise researchers of Hawaiian biogeography in general or researchers of the silversword radiation in particular—e.g. that the silversword alliance is monophyletic and younger than Kaua‘i, that one ancestral lineage founded the radiation, that they preferentially colonized younger islands in accordance with the progression rule—our framework greatly refines our ability to quantify exactly the location and timing of evolutionary events. This level of detail brings the next generation of biogeographic questions into reach: What geographical and ecological factors determine the periods of delay between island formation, island colonization, and radiation within the island? And how does the spatiotemporal distribution of habitat availability drive the evolution of novel ecological adaptations? By advancing the methodological framework to study these questions, we come closer to understanding the phenomenon of adaptive radiation as it behaves in nature.

The Hawaiian silversword alliance is representative of many island biogeographic systems: fossils are few or absent, island ages are uncertain, comprehensive genetic sampling is limited, but evolutionary hypotheses are plentiful. Recognizing these commonalities, we developed our inference strategy to be easily translated into other island biogeographic systems, even with systems that are not as well-behaved or well-understood as the Hawaiian silverswords. Our method, for instance, is directly relevant to the study of other Hawaiian flora and fauna, including the honeycreepers (Lerner et al. 2011), *Psychotria* (Nepokroeff et al. 2003), mints (Lindqvist and Albert 2002), lobelioids (Givnish et al. 2009), drosophilid flies (Lapoint et al. 2013), hyposmocomid moths (Haines et al. 2014), and many other remarkable clades. But, also, the relaxed rock paleogeographic model we introduced readily accommodates origin sequences for island systems far more complex than that of the Hawaiian Archipelago, such as the Galápagos Islands (Geist et al. 2014) or the Indo-Australian Archipelago (Lohman et al. 2011). Following the pioneering work of Sanmartín et al. (2008), a joint analysis that pools biogeographic evidence across multiple clades could, in principle, allow one to estimate otherwise uncertain paleogeographic features, such as area age, availability, and connectivity.

For many biogeographic systems, even minor amounts of phylogenetic, biogeographic, and paleogeographic uncertainty can obscure our intuition about historical events. In many cases, the perception of such uncertainty is daunting enough to prevent further investigation, because it clouds our sense of what is tractable: it becomes unclear what features of a clade’s history can be reconstructed and at what level of detail can hypotheses be posed and tested. What we show in this work is that by designing our inference methods to embrace these inherent sources of uncertainty, we can still form and test biogeographic hypotheses against a range of plausible, but ultimately unknowable, evolutionary histories, leading us to better understand how species diversify in space and time.

## 6 Contributions

M.J.L., W.A.F., and B.G.B. designed the study. M.J.L. and W.A.F. developed the methods and conducted the analyses. M.J.L, W.A.F., and B.G.B. interpreted the results and wrote the manuscript.

## 7 Acknowledgements

We thank Miranda Sinnott-Armstrong, Jun Ying Lim, Tracy Heath, and the Heath Lab for sharing insights that improved an early draft of this manuscript.

## References

Allison, P. A. and D. J. Bottjer. 2011. Taphonomy: bias and process through time. Pages 1–17 *in* Taphonomy (N. H. Landman and P. J. Harries, eds.). Springer, Heidelberg, Germany.

Baldwin, B. G. 1997. Adaptive radiation of the Hawaiian silversword alliance: congruence and conflict of phylogenetic evidence from molecular and non-molecular investigations. Pages 103–128 *in* Molecular evolution and adaptive radiation (T. J. Givnish and K. J. Sytsma, eds.). Cambridge University Press Cambridge, Cambridge, UK.

Baldwin, B. G. 2003a. A phylogenetic perspective on the origin and evolution of Madiinae. Pages 193–228 *in* Tarweeds & silverswords: evolution of the Madiinae (S. Carlquist, B. G. Baldwin, and G. D. Carr, eds.). Missouri Botanical Garden Press, St. Louis, Missouri, USA.

Baldwin, B. G. 2003b. Natural history of the continental tarweeds and the Hawaiian silversword alliance. Pages 1–16 *in* Tarweeds & silverswords: evolution of the Madiinae (S. Carlquist, B. G. Baldwin, and G. D. Carr, eds.). Missouri Botanical Garden Press, St. Louis, Missouri, USA.

Baldwin, B. G. 2014. Origins of plant diversity in the California Floristic Province. Annual Review of Ecology, Evolution, and Systematics 45:347–369.

Baldwin, B. G., D. W. Kyhos, and J. Dvorak. 1990. Chloroplast DNA evolution and adaptive radiation in the Hawaiian silversword alliance (Asteraceae-Madiinae). Annals of the Missouri Botanical Garden Pages 96–109.

Baldwin, B. G., D. W. Kyhos, J. Dvorak, and G. D. Carr. 1991. Chloroplast DNA evidence for a North American origin of the Hawaiian silversword alliance (Asteraceae). Proceedings of the National Academy of Sciences 88:1840–1843.

Baldwin, B. G. and R. H. Robichaux. 1995. Historical biogeography and ecology of the Hawaiian silversword alliance (Asteraceae): new molecular phylogenetic perspectives. Pages 259–287 *in* Hawaiian biogeography: evolution on a hot spot archipelago (W. L. Wagner and V. A. Funk, eds.). Smithsonian Institution Press, Washington, DC, USA.

Baldwin, B. G. and M. J. Sanderson. 1998. Age and rate of diversification of the hawaiian silversword alliance (compositae). Proceedings of the National Academy of Sciences 95:9402–9406.

Baldwin, B. G. and B. L. Wessa. 2000. Origin and relationships of the tarweed–silversword lineage (Compositae–Madiinae). American Journal of Botany 87:1890–1908.

Baldwin, B. G., B. L. Wessa, and J. L. Panero. 2002. Nuclear rDNA evidence for major lineages of helenioid Heliantheae (Compositae). Systematic Botany 27:161–198.

Barreda, V. D., L. Palazzesi, M. C. Tellería, E. B. Olivero, J. I. Raine, and F. Forest. 2015. Early evolution of the angiosperm clade Asteraceae in the Cretaceous of Antarctica. Proceedings of the National Academy of Sciences 112:10989–10994.

Blonder, B., B. G. Baldwin, B. J. Enquist, and R. H. Robichaux. 2016. Variation and macroevolution in leaf functional traits in the Hawaiian silversword alliance (Asteraceae). Journal of Ecology 104:219–228.

Burney, D. A., H. F. James, L. P. Burney, S. L. Olson, W. Kikuchi, W. L. Wagner, M. Burney, D. McCloskey, D. Kikuchi, F. V. Grady, et al. 2001. Fossil evidence for a diverse biota from Kaua’i and its transformation since human arrival. Ecological Monographs 71:615–641.

Carlquist, S. 1974. Island Biology. Columbia University Press: New York & London.

Carr, G. D. 1985. Monograph of the Hawaiian Madiinae (Asteraceae): *Argyroxiphium*, *Dubautia*, and *Wilkesia*. Allertonia 4:1–123.

Carr, G. D. 1999. A reassessment of *Dubautia* (Asteraceae: Heliantheae–Madiinae) on Kaua‘i. Pacific Science 53:144–158.

Carr, G. D. 2003. Chromosome evolution in Madiinae. Pages 53–78 *in* Tarweeds & silverswords: evolution of the Madiinae (S. Carlquist, B. G. Baldwin, and G. D. Carr, eds.). Missouri Botanical Garden Press, St. Louis, Missouri, USA.

Carr, G. D. and D. W. Kyhos. 1986. Adaptive radiation in the Hawaiian silversword alliance (Compositae–Madiinae). II. Cytogenetics of artificial and natural hybrids. Evolution 40:959–976.

Carson, H. L. and D. A. Clague. 1995. Geology and biogeography of the Hawaiian Islands. Smithsonian Institution Press Washington (DC).

Clague, D. A. and G. B. Dalrymple. 1987. The Hawaiian-Emperor volcanic chain. part I. Geologic evolution. Pages 5–54 *in* Volcanism in Hawaii (R. W. Decker, T. L. Wright, and P. H. Stauffer, eds.). US Geological Survey Professional Paper 1350, U.S. Government Printing Office, Washington, DC, USA.

Clague, D. A. and D. R. Sherrod. 2014. Growth and degradation of Hawaiian volcanoes vol. 1801. US Geol. Surv. Prof. Pap.

Cowie, R. H. and B. S. Holland. 2008. Molecular biogeography and diversification of the endemic terrestrial fauna of the Hawaiian Islands. Philosophical Transactions of the Royal Society of London B: Biological Sciences 363:3363–3376.

Drummond, A. J., S. Y. Ho, M. J. Phillips, and A. Rambaut. 2006. Relaxed phylogenetics and dating with confidence. PLoS Biology 4:e88.

Dupin, J., N. J. Matzke, T. Särkinen, S. Knapp, R. G. Olmstead, L. Bohs, and S. D. Smith. 2017. Bayesian estimation of the global biogeographical history of the Solanaceae. Journal of Biogeography 44:887–899.

Freyman, W. A. and S. Höhna. 2017. Cladogenetic and anagenetic models of chromosome number evolution: a Bayesian model averaging approach. Systematic Biology syx065.

Funk, V. A. and W. L. Wagner. 1995. Biogeographic patterns in the hawaiian islands. Pages 379–419 *in* Hawaiian biogeography: evolution on a hot spot archipelago (W. L. Wagner and V. A. Funk, eds.). Smithsonian Institution Press, Washington, DC, USA.

Gavrilets, S. and J. B. Losos. 2009. Adaptive radiation: contrasting theory with data. Science 323:732–737.

Geist, D. J., H. Snell, H. Snell, C. Goddard, and M. D. Kurz. 2014. A paleogeographic model of the Galápagos Islands and biogeographical and evolutionary implications. The Galápagos: a natural laboratory for the Earth Sciences. American Geophysical Union, Washington DC, USA Pages 145–166.

Gillespie, R. 2004. Community assembly through adaptive radiation in Hawaiian spiders. Science 303:356–359.

Gillespie, R. G., E. M. Claridge, and S. L. Goodacre. 2008. Biogeography of the fauna of French Polynesia: diversification within and between a series of hot spot archipelagos. Philosophical Transactions of the Royal Society of London B: Biological Sciences 363:3335–3346.

Gillespie, R. G., H. B. Croom, and G. L. Hasty. 1997. Phylogenetic relationships and adaptive shifts among major clades of *Tetragnatha* spiders (araneae: Tetragnathidae) in hawai‘i. Pacific Science 51:380–394.

Givnish, T. J. 2015. Adaptive radiation versus ‘radiation’ and ‘explosive diversification’: why conceptual distinctions are fundamental to understanding evolution. New Phytologist 207:297–303.

Givnish, T. J., K. C. Millam, A. R. Mast, T. B. Paterson, T. J. Theim, A. L. Hipp, J. M. Henss, J. F. Smith, K. R. Wood, and K. J. Sytsma. 2009. Origin, adaptive radiation and diversification of the Hawaiian lobeliads (Asterales: Campanulaceae). Proceedings of the Royal Society of London B: Biological Sciences 276:407–416.

Goldberg, E. E. and B. Igić. 2012. Tempo and mode in plant breeding system evolution. Evolution 66:3701–3709.

Goldberg, E. E., L. T. Lancaster, and R. H. Ree. 2011. Phylogenetic inference of reciprocal effects between geographic range evolution and diversification. Systematic Biology 60:451–465.

Grant, P. R., B. R. Grant, J. A. Markert, L. F. Keller, and K. Petren. 2004. Convergent evolution of Darwin’s finches caused by introgressive hybridization and selection. Evolution 58:1588–1599.

Haines, W. P., P. Schmitz, and D. Rubinoff. 2014. Ancient diversification of Hyposmocoma moths in Hawaii. Nature Communications 5:3502.

Hasegawa, M., H. Kishino, and T. Yano. 1985. Dating the human-ape splitting by a molecular clock of mitochondrial DNA. Journal of Molecular Evolution 22:160–174.

Heath, T. A., J. P. Huelsenbeck, and T. Stadler. 2014. The fossilized birth–death process for coherent calibration of divergence-time estimates. Proceedings of the National Academy of Sciences 111:E2957–E2966.

Höhna, S., M. J. Landis, T. A. Heath, B. Boussau, N. Lartillot, B. R. Moore, J. P. Huelsenbeck, and F. Ronquist. 2016. RevBayes: Bayesian phylogenetic inference using graphical models and an interactive model-specification language. Systematic Biology 65:726–736.

Hotchkiss, S. and J. O. Juvik. 1999. A Late-Quaternary pollen record from Ka‘au Crater, O‘ahu, Hawai‘i. Quaternary Research 52:115–128.

Jacobs, D. K., T. A. Haney, and K. D. Louie. 2004. Genes, diversity, and geologic process on the Pacific coast. Annual Review of Earth and Planetary Sciences 32:601–652.

Judd, W. S., C. S. Campbell, E. Kellogg, P. Stevens, and M. Donoghue. 2016. Plant systematics: a phylogenetic approach. Oxford University Press, Oxford, UK.

Kamath, A. and J. B. Losos. 2017. Does ecological specialization transcend scale? Habitat partitioning among individuals and species of *Anolis* lizards. Evolution 71:541–549.

Lamichhaney, S., F. Han, J. Berglund, C. Wang, M. S. Almén, M. T. Webster, B. R. Grant, P. R. Grant, and L. Andersson. 2016. A beak size locus in Darwin’s finches facilitated character displacement during a drought. Science 352:470–474.

Landis, M. J. 2017. Biogeographic dating of speciation times using paleogeographically informed processes. Systematic Biology 66:128–144.

Landis, M. J., N. J. Matzke, B. R. Moore, and J. P. Huelsenbeck. 2013. Bayesian analysis of biogeography when the number of areas is large. Systematic Biology 62:789–804.

Lapoint, R. T., P. M. O’Grady, and N. K. Whiteman. 2013. Diversification and dispersal of the Hawaiian Drosophilidae: The evolution of Scaptomyza. Molecular Phylogenetics and Evolution 69:95–108.

Lerner, H. R., M. Meyer, H. F. James, M. Hofreiter, and R. C. Fleischer. 2011. Multilocus resolution of phylogeny and timescale in the extant adaptive radiation of Hawaiian honeycreepers. Current Biology 21:1838–1844.

Lim, J. Y. and C. R. Marshall. 2017. The true tempo of evolutionary radiation and decline revealed on the Hawaiian archipelago. Nature 543:710–713.

Lindqvist, C. and V. A. Albert. 2002. Origin of the Hawaiian endemic mints within North American Stachys (Lamiaceae). American Journal of Botany 89:1709–1724.

Lohman, D. J., M. de Bruyn, T. Page, K. von Rintelen, R. Hall, P. K. L. Ng, H.-T. Shih, G. R. Carvalho, and T. von Rintelen. 2011. Biogeography of the Indo-Australian archipelago. Annual Review of Ecology, Evolution, and Systematics 42:205–226.

Losos, J. B. 1992. The evolution of convergent structure in Caribbean *Anolis* communities. Systematic Biology 41:403–420.

Losos, J. B., T. R. Jackman, A. Larson, D. de Queiroz, and L. Rodríguez-Schettino. 1998a. Contingency and determinism in replicated adaptive radiations of island lizards. Science 279:2115–2118.

Losos, J. B., T. R. Jackman, A. Larson, K. de Queiroz, and L. Rodríguez-Schettino. 1998b. Contingency and determinism in replicated adaptive radiations of island lizards. Science 279:2115–2118.

Mahler, D. L., T. Ingram, L. J. Revell, and J. B. Losos. 2013. Exceptional convergence on the macroevolutionary landscape in island lizard radiations. Science 341:292–295.

Massana, K. A., J. M. Beaulieu, N. J. Matzke, and B. C. O’Meara. 2015. Non-null Effects of the Null Range in Biogeographic Models: Exploring Parameter Estimation in the DEC Model. bioRxiv Page 026914.

Nee, S., R. M. May, and P. H. Harvey. 1994. The reconstructed evolutionary process. Philosophical Transactions of the Royal Society of London B 344:305–311.

Nepokroeff, M., K. J. Sytsma, W. L. Wagner, and E. A. Zimmer. 2003. Reconstructing ancestral patterns of colonization and dispersal in the Hawaiian understory tree genus Psychotria (Rubiaceae): a comparison of parsimony and likelihood approaches. Systematic Biology 52:820–838.

Nielsen, R. 2002. Mapping mutations on phylogenies. Systematic Biology 51:729–739.

Olson, S. L. and H. F. James. 1982. Prodromus of the fossil avifauna of the Hawaiian Islands. Smithsonian Institution Press, Washington, DC, USA.

Osborn, H. F. 1902. The law of adaptive radiation. The American Naturalist 36:353–363.

Parent, C. E., A. Caccone, and K. Petren. 2008. Colonization and diversification of Galápagos terrestrial fauna: a phylogenetic and biogeographical synthesis. Philosophical Transactions of the Royal Society of London B: Biological Sciences 363:3347–3361.

Poe, S., A. Nieto-montes de oca, O. Torres-carvajal, K. De Queiroz, J. A. Velasco, B. Truett, L. N. Gray, M. J. Ryan, G. Köhler, F. Ayala-varela, et al. 2017. A phylogenetic, biogeographic, and taxonomic study of all extant species of *Anolis* (Squamata; Iguanidae). Systematic Biology 66:663–697.

Price, J. P. and D. A. Clague. 2002. How old is the Hawaiian biota? Geology and phylogeny suggest recent divergence. Proceedings of the Royal Society of London B: Biological Sciences 269:2429–2435.

Raven, P. H. and D. I. Axelrod. 1978. Origin and relationships of the California flora vol. 72. University of California Press, Berkeley, CA, USA.

Ree, R. H., B. R. Moore, C. O. Webb, and M. J. Donoghue. 2005. A likelihood framework for inferring the evolution of geographic range on phylogenetic trees. Evolution 59:2299–2311.

Ree, R. H. and S. A. Smith. 2008. Maximum likelihood inference of geographic range evolution by dispersal, local extinction, and cladogenesis. Systematic Biology 57:4–14.

Rodrigue, N., H. Philippe, and N. Lartillot. 2008. Uniformization for sampling realizations of Markov processes: Applications to Bayesian implementations of codon substitution models. Bioinformatics 24:56–62.

Sakai, A. K., W. L. Wagner, and L. A. Mehrhoff. 2002. Patterns of endangerment in the Hawaiian flora. Systematic Biology 51:276–302.

Sanmartín, I., P. V. D. Mark, and F. Ronquist. 2008. Inferring dispersal: a bayesian approach to phylogeny-based island biogeography, with special reference to the canary islands. Journal of Biogeography 35:428–449.

Sansom, R. S., S. E. Gabbott, and M. A. Purnell. 2010. Non-random decay of chordate characters causes bias in fossil interpretation. Nature 463:797–800.

Sato, A., H. Tichy, C. O’hUigin, P. R. Grant, B. R. Grant, and J. Klein. 2001. On the origin of Darwin’s finches. Molecular Biology and Evolution 18:299–311.

Schluter, D. 2000. The ecology of adaptive radiation. Oxford University Press, New York, New York, USA.

Shaw, K. L. and R. G. Gillespie. 2016. Comparative phylogeography of oceanic archipelagos: Hotspots for inferences of evolutionary process. Proceedings of the National Academy of Sciences 113:7986–7993.

Silvertown, J., J. Francisco-Ortega, and M. Carine. 2005. The monophyly of island radiations: an evaluation of niche pre-emption and some alternative explanations. Journal of Ecology 93:653–657.

Simpson, G. G. 1944. Tempo and mode in evolution. Columbia University Press, New York, USA.

Thorne, J., H. Kishino, and I. S. Painter. 1998. Estimating the rate of evolution of the rate of molecular evolution. Molecular Biology and Evolution 15:1647–1657.

Wagner, W. L., D. R. Herbst, and D. H. Lorence. 2005. Flora of the Hawaiian Islands website. http://botany.si.edu/pacificislandbiodiversity/hawaii Accessed: 2017-08-25.

Wagner, W. L., S. Weller, and A. Sakai. 1995. Phylogeny and biogeography in schiedea and alsinidendron (caryophyllaceae). Pages 221–258 *in* Hawaiian biogeography: evolution on a hot spot archipelago (W. L. Wagner and V. A. Funk, eds.). Smithsonian Institution Press, Washington, DC, USA.

Webb, C. O. and R. H. Ree. 2012. Historical biogeography inference in Malesia. Pages 191–215 *in* Biotic evolution and environmental change in Southeast Asia (D. Gower, K. Johnson, J. Richardson, B. Rosen, L. Ruber, and S. Williams, eds.) Cambridge University Press.

Whittaker, R. J., K. A. Triantis, and R. J. Ladle. 2008. A general dynamic theory of oceanic island biogeography. Journal of Biogeography 35:977–994.

Witter, M. S. and G. D. Carr. 1988. Adaptive radiation and genetic differentiation in the Hawaiian silversword alliance (Compositae: Madiinae). Evolution 42:1278–1287.

Yang, Z., N. Goldman, and A. E. Friday. 1995a. Maximum likelihood trees from DNA sequences: A peculiar statistical estimation problem. Systematic Biology 44:384–399.

Yang, Z., S. Kumar, and M. Nei. 1995b. A new method for inference of ancestral nucleotide and amino acid sequences. Genetics 141:1641–1650.

Zuckerkandl, E. and L. Pauling. 1962. Molecular disease, evolution, and genetic heterogeneity. Pages 189–225 *in* Horizons in Biochemistry (M. Kasha and B. Pullman, eds.) Academic Press, New York.

